# *APOE3* Christchurch modulates tau phosphorylation and β-catenin/Wnt/Cadherin signaling in induced pluripotent stem cell-derived cerebral organoids from Alzheimer’s cases

**DOI:** 10.1101/2023.01.11.523290

**Authors:** RC Mazzarino, P Perez-Corredor, TE Vanderleest, GN Vacano, JS Sanchez, ND Villalba-Moreno, S Krausemann, MA Mendivil-Perez, D Aguillón, M Jimenez-Del-Río, A Baena, D Sepulveda-Falla, FJ Lopera, YT Quiroz, JF Arboleda-Velasquez

## Abstract

Alzheimer’s disease (AD) is the most common cause of dementia among older adults. *APOE3* Christchurch (R136S, *APOE3Ch*) variant homozygosity was reported in an individual with extreme resistance to autosomal dominant AD due to the *PSEN1* E280A mutation. This subject had a delayed clinical age at onset and resistance to tauopathy and neurodegeneration despite extremely high amyloid plaque burden. We established induced pluripotent stem (iPS) cell-derived cerebral organoids from this resistant case and from a non-protected kindred control (with *PSEN1* E280A and *APOE3/3*). We used CRISPR/Cas9 gene editing to successfully remove the *APOE3Ch* to wild type in iPS cells from the protected case and to introduce the *APOE3Ch* as homozygote in iPS cells from the non-protected case to examine causality. We found significant reduction of tau phosphorylation (pTau 202/205 and pTau396) in cerebral organoids with the *APOE3Ch* variant, consistent with the strikingly reduced tau pathology found in the resistant case. We identified Cadherin and Wnt pathways as signaling mechanisms regulated by the *APOE3Ch* variant through single cell RNA sequencing in cerebral organoids. We also identified elevated β-catenin protein, a regulator of tau phosphorylation, as a candidate mediator of *APOE3Ch* resistance to tauopathy. Our findings show that *APOE3Ch* is necessary and sufficient to confer resistance to tauopathy in an experimental *ex-vivo* model establishing a foundation for the development of novel, protected case-inspired therapeutics for tauopathies, including Alzheimer’s.

## Introduction

Alzheimer’s disease (AD) is the most common cause of dementia among older adults. AD affects an estimated 55 million people worldwide with numbers expected to exceed 152 million people by the year 2050 (Patterson 2018). AD is characterized by the formation of amyloid plaques and tau fibrils in the brain as well as calcium and mitochondrial dysregulation that manifests in neuronal death and memory deficits (Knopman et al. 2021; Nunomura et al. 2001; Mosconi 2013). Autosomal Dominant Alzheimer’s disease (ADAD) accounts for approximately 1% of diagnosed patients (Sims, Hill, and Williams 2020), with about 70% of ADAD patients having a Presenilin 1 (*PSEN1*) mutation (Sun et al. 2017). Recently, a member of the Colombian *PSEN1* E280A (Paisa) kindred was identified as being resistant to ADAD. Carriers of the *PSEN1* E280A mutation develop mild cognitive impairment at 43-45 and dementia at 49-50 years of age (95% confidence intervals); the identified female patient did not develop mild cognitive impairment until her seventies. She had very limited levels of tau pathology, neuroinflammation, and neurodegeneration, but extremely high levels of amyloid plaque burden (Arboleda-Velasquez et al. 2019; Sepulveda-Falla et al. 2022).

She was also found to be homozygous for the Apolipoprotein E3 Christchurch (*APOE3Ch*) variant (R136S) which was identified as a candidate variant responsible for her resistance to ADAD (Arboleda-Velasquez et al. 2019). Genetic imputation of causality could not be confirmed because only a single *APOE3Ch* homozygote case with resistance was identified. In this study, we used cells from resistant and non-resistant patients to generate iPS cells, used genomic editing to introduce or remove the *APOE3Ch* or *PSEN1* E280A mutations and identified the Cadherin/Wnt/β-catenin signaling pathways as plausible components by which the *APOE3Ch* mutation acts to provide resistance to ADAD.

## Results

### ADAD cases selection

We identified two informative patients for this study (named α and ω to protect privacy): Patient α was previously described as the homozygote *APOE3Ch* protected case (Arboleda-Velasquez et al. 2019) and Patient ω, not previously described, was selected as a control for this study. Both individuals are from the Paisa *PSEN1* E280A kindred and female, though not closely related. Patient α developed mild cognitive impairment (MCI) in her seventies and was seen as protected against ADAD while Patient ω developed MCI and dementia in her forties as expected for this kindred. PET imaging of each patient was performed (Figure 1) to evaluate amyloid and tau burden. Pittsburgh compound B PET imaging for amyloid plaques revealed elevated burden in Patient α over Patient ω while [18F]Flortaucipir PET imaging displayed significant burden in the medial temporal and parietal regions of Patient ω, which is markedly reduced in Patient α. Taken together, these individuals were selected as an informative pairing for protected and non-protected cases.

**Figure 1:**
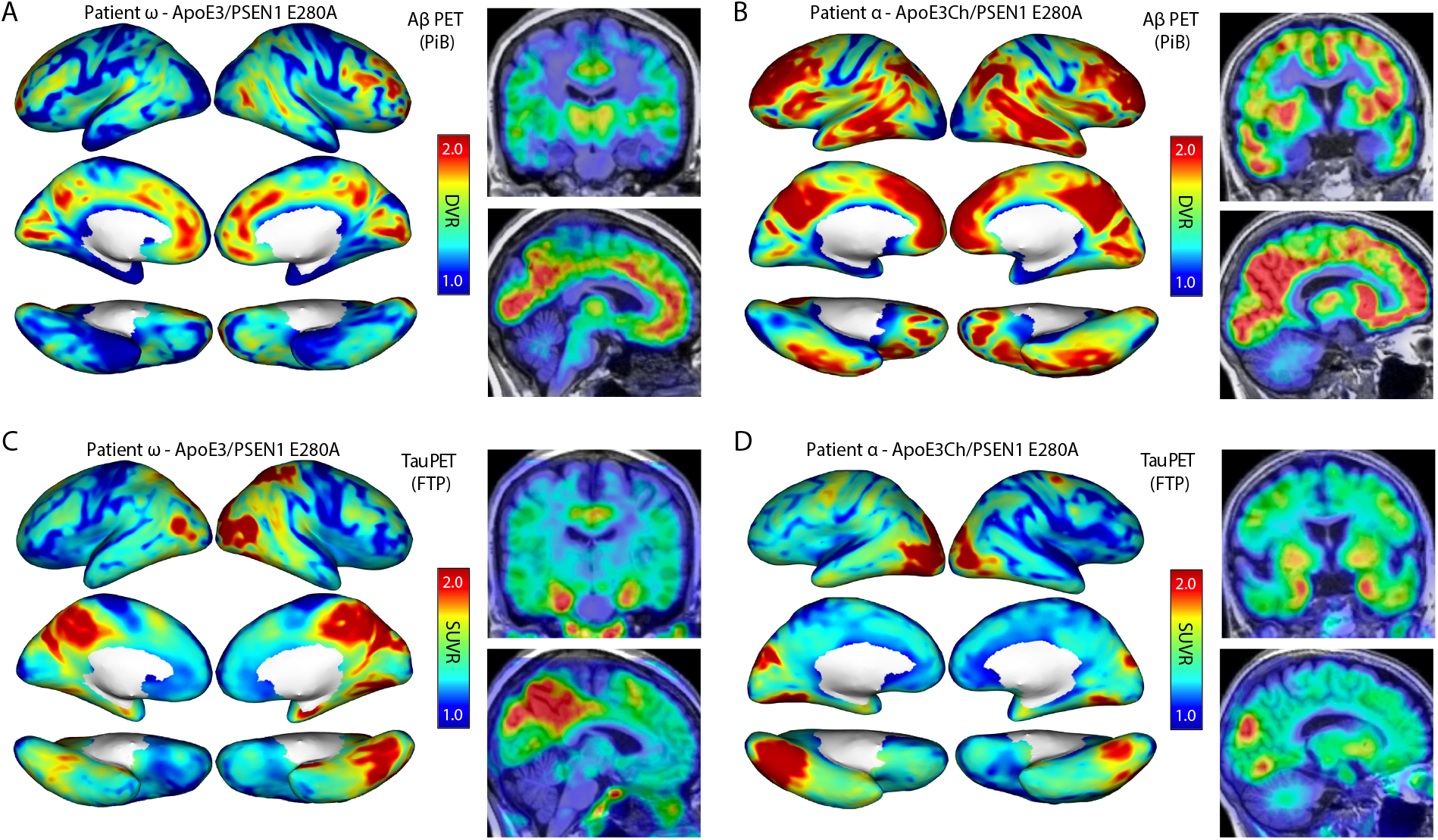
Patient *in vivo* neuroimaging. Patient ω is a non-protected, Paisa kindred control patient and was scanned for Amyloid β **(A)** and tau burden **(C)** Patient α is a protected patient identified in the was previously described (scans previously used in (Arboleda-Velasquez et al. 2019)) and scanned for Amyloid β **(B)** and tau burden **(D)**. We have identified two specific patients for this study. Patient α was previously described (Arboleda-Velasquez et al. 2019) and Patient ω is novel for this study. Both individuals are within the Paisa PSEN1 E280A kindred and female. Scans indicated elevated Amyloid β burden in Patient α **(B)** compared to Patient ω **(A)**. Tau burden was elevated in Patient ω **(C)** over Patient α **(D)**, with marked increase in medial temporal and parietal regions. PiB=Pittsburgh Compound B; FTP=Flortaucipir; DVR=distribution volume ratio; SUVR=standardized uptake value ratio.

### Generation and gene editing of patient-derived induced pluripotent stem cells

*We* generated patient-derived iPS cell lines to examine the effects of the *APOE3Ch* R136S as well as *PSEN1* E280A mutation (Figure 2A). Patient blood samples were reprogramed by Sendai virus containing Yamanaka factors (*OCT4, SOX2, KLF4, and c-MYC*), subcloned, and tested by immunofluorescence (IF) imaging for the pluripotency markers SSEA4, Oct4, Tra-1-60, and Nanog (Figure 2B). Reprogrammed subclones were then edited using CRISPR Cas9 to knock in gene variants. From Patient α <u>with the *PSEN1* E280A mutation, *APOE3* homozygosity, and extreme resistance to Alzheimer’s we obtained the following cell lines:</u> 1) we used CRISPR to change the *APOE3Ch* to *APOE3* wild type (removal of putative protective factor, genotype= *PSEN1* E280A; *APOE3* WT); 2) we used CRISPR to correct the *PSEN1* E280A to *PSEN1* wild type (removal of Alzheimer’s causality factor, genotype = *PSEN1* WT; *APOE3Ch* homozygous); 3) unsuccessful gene editing control from *PSEN1* targeting; and 4) unsuccessful gene editing control from *APOE3* targeting. These unsuccessful gene editing controls represent isogenic parental controls of Patient α base genotype, *APOE3Ch* and *PSEN1* E280A. From Patient ω <u>with *PSEN1* E280A mutation and expected age of onset (non-protected individual) we obtained the following cell lines:</u> 1) We used CRISPR to correct the *PSEN1* E280A mutation to *PSEN1* wild type (removal of Alzheimer’s causality factor, genotype= *PSEN1* WT; *APOE3* WT); 2) We used CRISPR to introduce the *APOE3Ch* as homozygote (introduction of putative protective factor, genotype= *PSEN1* E280A; *APOE3Ch* homozygous); 3) unsuccessful gene editing control from *PSEN1* targeting; and 4) unsuccessful gene editing control from *APOE3* targeting. These unsuccessful gene editing controls represent isogenic parental controls of Patient ω base genotype of *APOE3* and *PSEN1* E280A. (Figure 2A)

**Figure 2:**
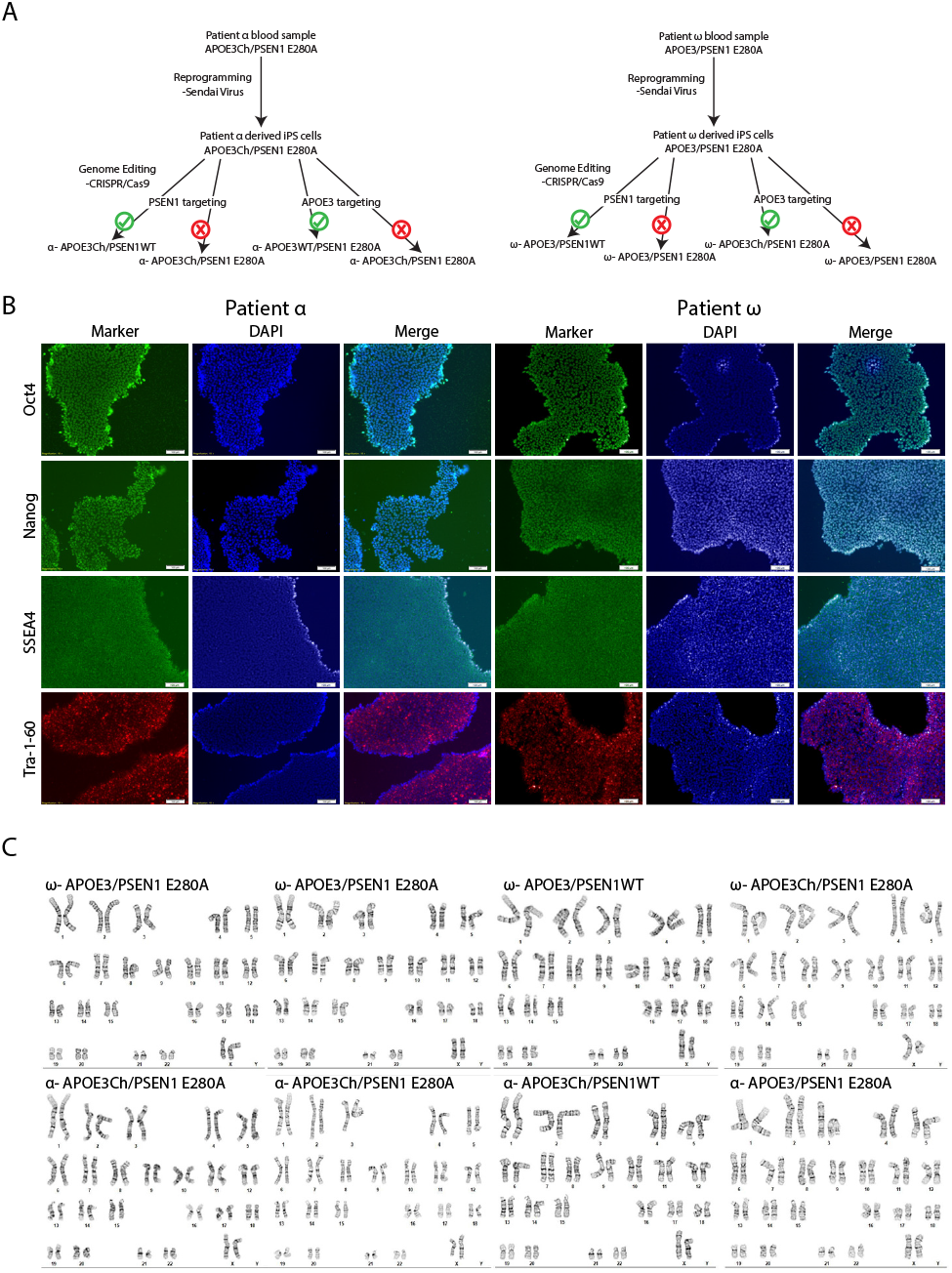
iPS cell line basic characterization. Patient derived iPS cell lines were designed to examine the effects of *PSEN1* E280A mutation as well as the *APOE3Ch* mutation **(A)**. Patient blood samples were reprogramed, subcloned, and tested by IF imaging for the pluripotency markers SSEA4, Oct4, Tra-1-60, and Nanog, scale bars 100 μm **(B)**. Karyotyping was performed and all cell lines generated showed 46 chromosomes and no overt abnormalities **(C)**.

Sanger sequencing was performed to confirm genetic edit success and isogenic controls were also selected from cells with unsuccessful gene editing that underwent all processes in parallel. Altogether, eight cell lines were generated for this study (four from each case). Karyotyping was performed and all cell lines generated showed 46 chromosomes without overt abnormalities (Figure 2C). Cell lines showed success for trilineage; embryoid bodies were formed and pushed towards ectoderm, mesoderm, and endoderm for two weeks and assessed by a three gene qPCR panel: Ectoderm-EN1, MAP2, NR2F2, Mesoderm-SNAIL2, RGS4, HAND2, Endoderm-SST, Klf5, AFP (data not shown).

### Differentiation of patient derived iPS into cerebral organoids

We chose a Lancaster type differentiation protocol offering a more basic, generalized cerebral organoid to identify processes that are based on cell types found within a central nervous system rather than in enriched population of specific cerebral cell type. No notable differences were observed between cell lines in the iPS phase initially and they grew as expected in mTeSR Plus media. All cell lines examined here showed positive progression during spheroid formation (5 days), cerebral organoid induction (2 days), expansion (3 days), and maturation (19 days) in accordance with manufacturer’s protocol timeline. Organoids were fixed, paraffin embedded, and then sections were stained for cell type markers. Immunohistochemistry (IHC) for cell type markers showed positive staining for neurons (MAP2), early neurons/neural progenitors (PAX6), and early astrocytes in all organoids (Figure 3A, B). Minimal staining was noted for oligodendrocytes (Olig2). Organoids were negative for the mature astrocyte marker GFAP and the microglial marker IBA1 (Figure 3A). These results were expected with the differentiation protocol used and time points selected.

**Figure 3:**
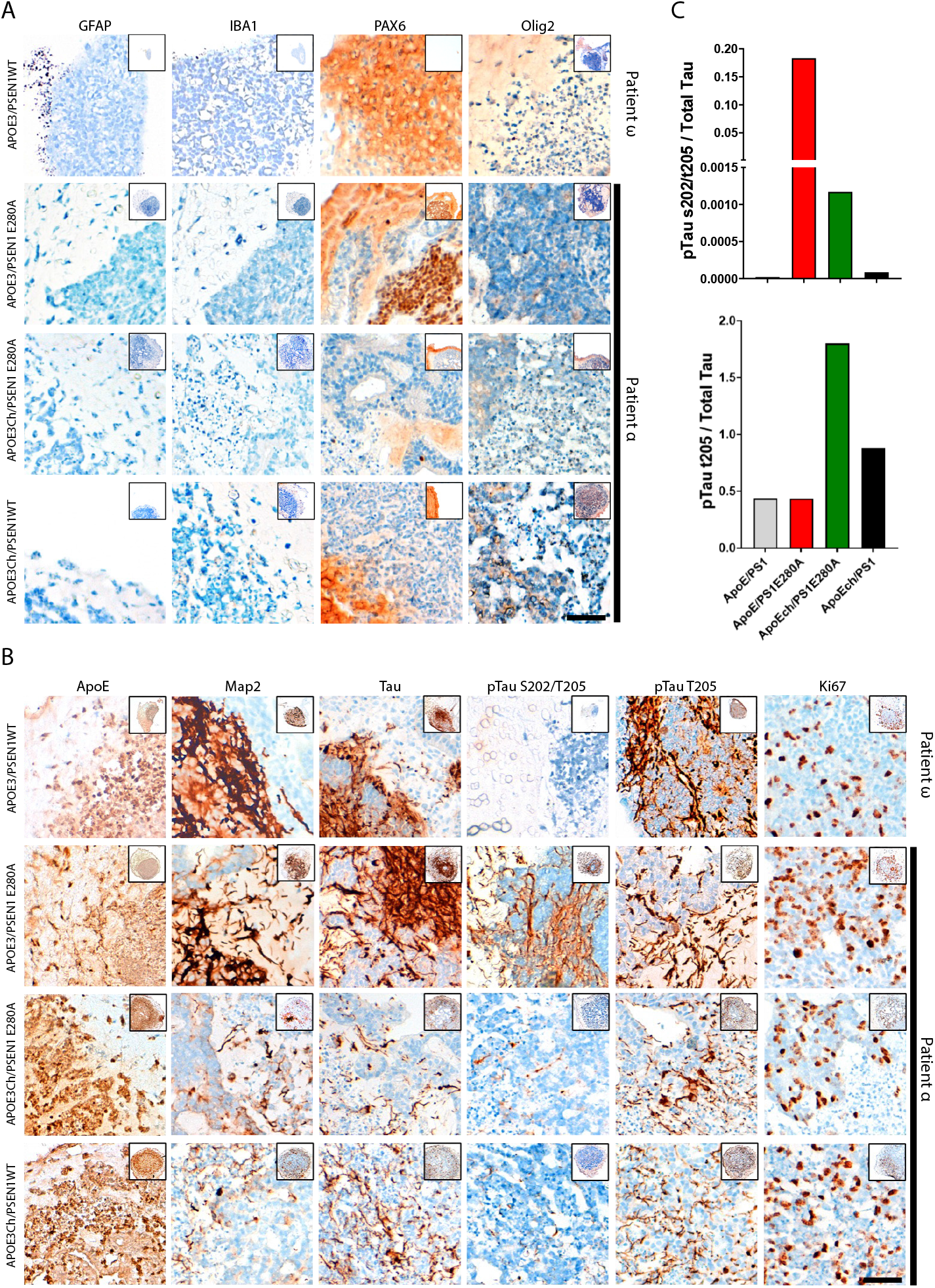
Histological and morphological analysis of *PSEN1* E280A *APOE3Ch* brain organoids and isogenic controls. Representative images of immunohistochemistry for GFAP (glia fibrillary acid protein), IBA1 (ionized calcium binding adapter 1), PAX6 (paired box protein 6) and Olig2 (oligodendrocyte transcription factor) **(A)**. Representative images of immunohistochemistry for ApoE, MAP2 (microtubule associated protein 2), tau, pTau s202/t205 (hyperphosphorylated tau in serine 202 and threonine 205), pTau t205 (hyperphorolyted tau in threonine 205) and Ki67 (proliferation marker Ki-67) of same organoids from panel A **(B)**. Stainings were performed in cerebral organoids derived from a *APOE3 PSEN1* E280A carrier (Patient ω) in which the *PSEN1* E280A mutation was removed (ApoE/PS1), and from a homozygous *APOE3ChPSEN1* E280A carrier (Patient α) in which the *APOE3Ch* mutation was removed (*APOE*/PS1E280A), was not removed (*APOE3Ch*/PS1E280A) and only the *PSEN1* mutation was removed (*APOE3Ch*/PS1). Bar graphs representing signal intensity ratios of pTau s202/t205 and pTau t205 over total tau **(C)**. The cerebral organoids derived from Patient α and with the *APOE3Ch* mutation removed showed considerable higher pTau s202/t205 levels (see notched axis). Meanwhile, pTau t205 signal tended to be higher in organoids carrying the *APOE3Ch* mutation regardless of PSEN1 background. Bars = 100 mm.

### *APOE3Ch* cerebral organoids have a protective pattern of tau phosphorylation

We focused on the histological analysis of tau expression and phosphorylation because resistance to tauopathy was a neuropathological hallmark in the *APOE3Ch* homozygote case (Patient α), first observed by PET imaging and later confirmed postmortem (Arboleda-Velasquez et al. 2019; Sepulveda-Falla et al. 2022). AT8 pTau S202/T205 staining of paired helical filaments (PHF) in organoids derived from the protected case *(APOE3Ch/PSEN1* E280A) was lower compared to organoids derived from isogenic cells in which the *APOE3Ch* was removed *(APOE3/PSEN1* E280A). pTau S202/T205 staining was similar in organoids where the *PSEN1* mutation had been corrected compared to organoids from the parental line. Overall, these findings suggest that pathological tau phosphorylation induced by the *PSEN1* E280A mutation is reduced by the presence of *APOE3Ch* in our model. To complement these results, we examined tau phosphorylation at pTau T205, a site associated with protective effects of mitogen-activated protein (MAP) kinase p38γ (Ittner et al. 2016; 2020). We found increased staining of pTau T205 in organoids derived from cells with the *APOE3Ch* compared to controls, independent of the *PSEN1* genotype in our organoid model. Overall, we concluded that the *APOE3Ch* variant was necessary to produce a protective pattern of tau phosphorylation defined as low pTau S202/T205 and high pTau T205 in cerebral organoids derived from an AD-protected case (Figure 3B, C).

We conducted immunofluorescence analyses to confirm a genetic link between specific patterns of tau phosphorylation and *APOE3Ch* using anti-pTau S396 in organoids derived cells of the non-protected case. pTau S396 is an early marker of pathologic tau phosphorylation (Mondragón-Rodríguez et al. 2014). pTau S396 staining was lower in organoids from the nonprotected Patient ω in which the *APOE3Ch* was introduced compared to isogenic controls. pTau S396 staining was similar in organoids with the *PSEN1* E280A mutation and those corrected to *PSEN1WT* (Figure 4 A and B). Again, changes in pTau S396 did not depend on the *PSEN1* genotype. This staining pattern was consistent in Patient α, whereby removal of the *APOE3Ch* variant led to a significant increase in pTau S396 staining compared to all other Patient α cell lines (Figure 4 C and D). We concluded that the *APOE3Ch* variant was necessary and sufficient to produce a protective pattern of tau phosphorylation defined as low pTau S396 in cerebral organoids derived from an AD-protected and non-protected patient. Overall, the effects of *APOE* genotypes on tau phosphorylation were independent from *PSEN1* genotypes suggesting a direct effect of *APOE* genotypes on the status of tau phosphorylation.

**Figure 4:**
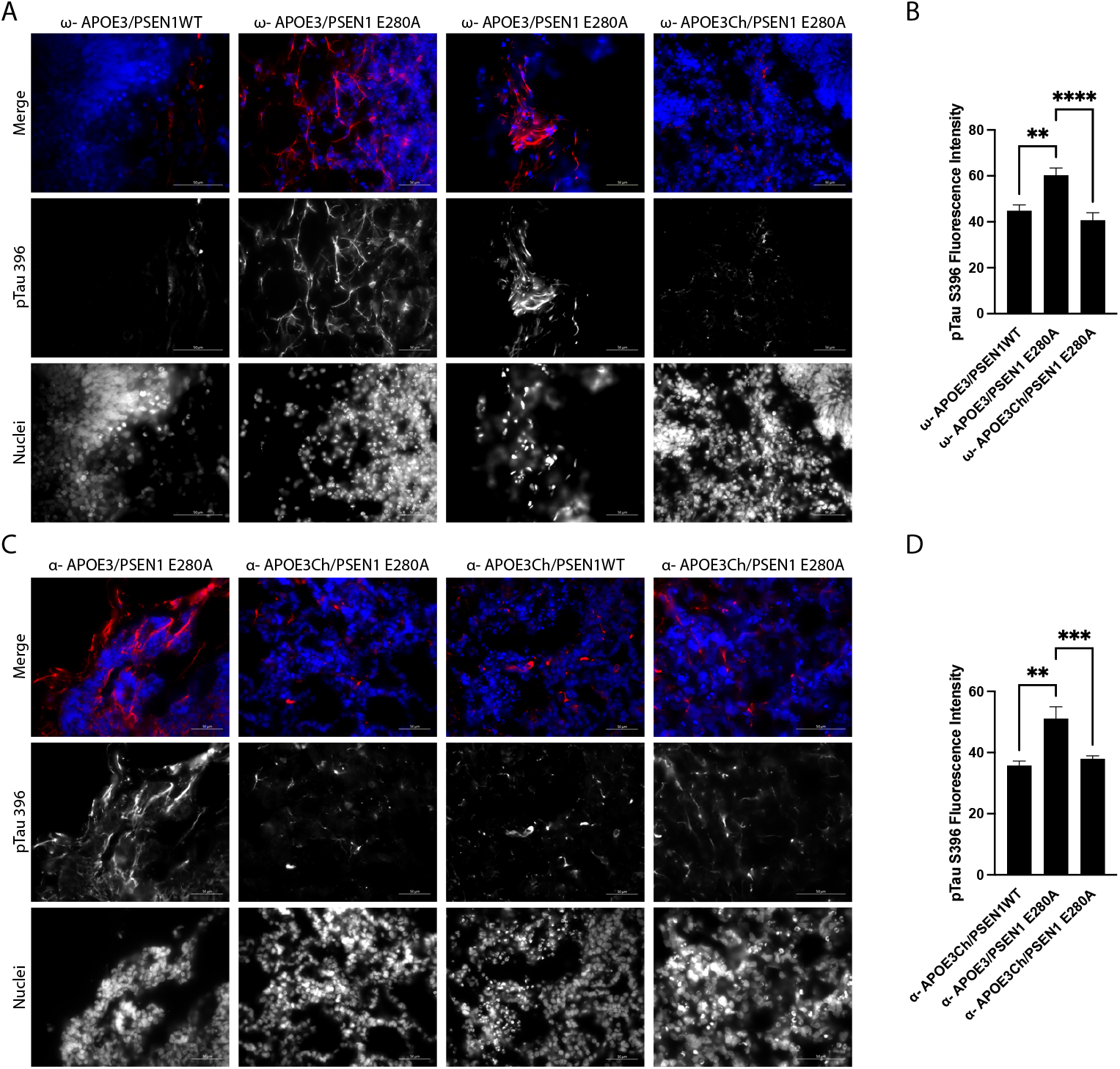
*APOE3Ch* decreases pTau S396. Cerebral organoids were formed and stained for nuclei and pTau S396 and imaged at 63X. Representative images from Patient ω **(A)** and Patient α **(C)** depicted, scale bars 50 μm. pTau S396 signal intensity was measured and isogenic controls for each patient were averaged together **(B, D).**

### Single cell RNA sequencing of cerebral organoids

We conducted scRNA-sequencing to examine potential mechanistic links between *APOE* genotypes and tau modification. Six organoids were randomly collected from each set of organoids with specific genotypes, dissociated, and cDNA libraries built using the 10X Genomics platform and Illumina sequenced. FASTQ files were initially processed through the 10X Genomics Cell Ranger suite and H5 output files were further processed through Seurat. Quality of datasets was confirmed within Seurat using standard QC measures (data available on request). We used DoubletFinder to reduce multiplet noise within our datasets. To be able to compare results within and between all cell lines, we downsampled all datasets to 6,500 cells then integrated all scRNA-seq output for all cell lines using Harmony and SCTransform. UMAPs for both patients exhibit overlap of cluster and cell types indicating that integration was successful (Figure 5A). By using GSEA C8 Cell Type Signature Gene Sets we were able to identify cell types (Figure 5B) and grouped like-clusters together forming superclusters (Figure 5C).

**Figure 5:**
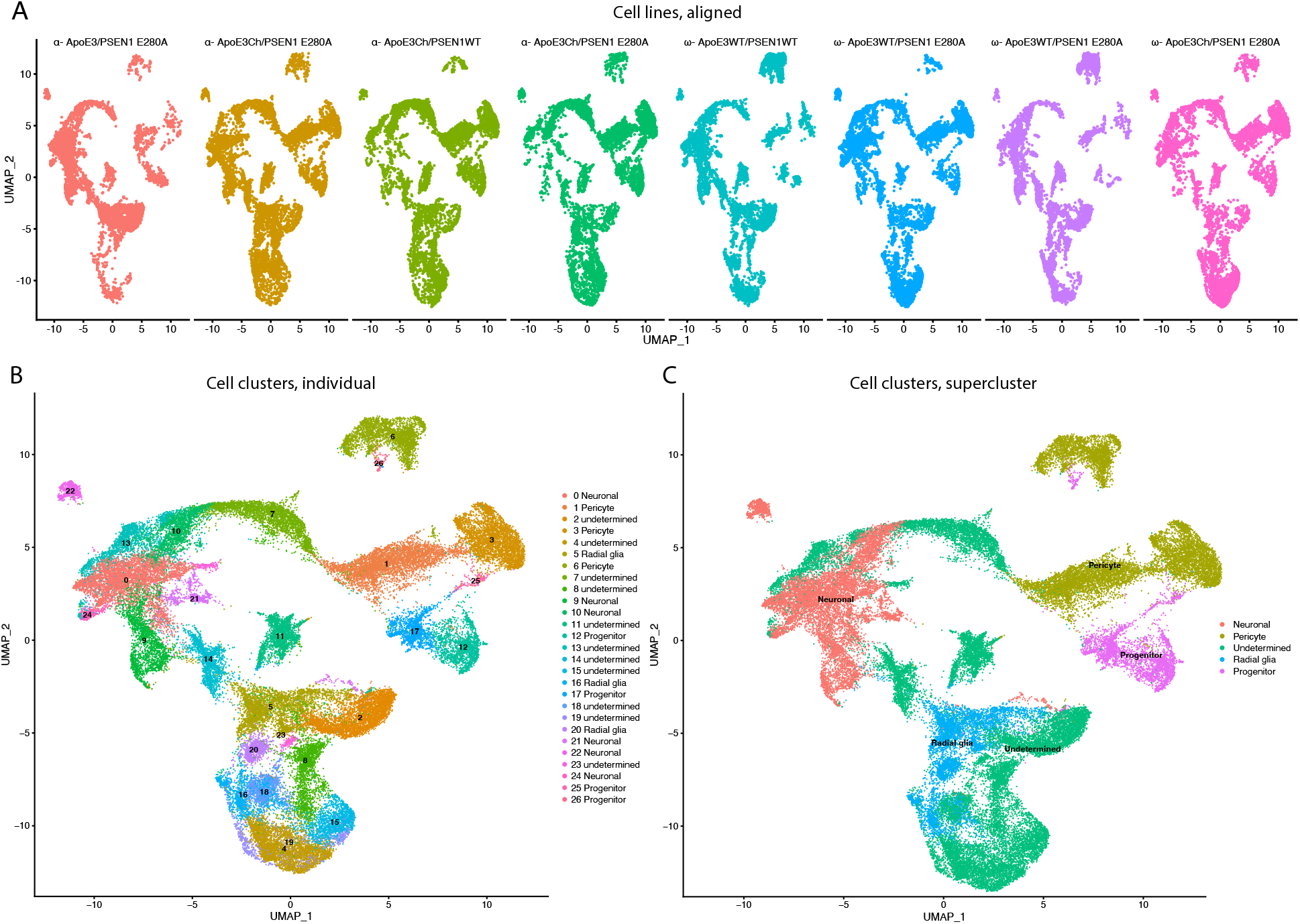
Cell line scRNAseq UMAP clusters characteristics. scRNAseq data was downsampled and integrated using SCTransform and Harmony. Individual cell line UMAPs show overlap indicating that integration was successful in approximating cell **(A)**. Cell cluster transcript lists were run through GSEA C8 Cell Type for cluster identity **(B)**. Similar cluster identities were combined to form superclusters **(C)**.

We reduced data dimensionality and converted the scRNA-seq data sets to pseudo-bulk to identify broad changes induced by the *APOE3Ch* variant. This was done to identify large differences across the datasets and to streamline workflow due to the number of cell lines tested and complexity within integrated datasets. We used the pseudo-bulk datasets to generate log2(FC >1 and <-1) differential expression gene lists of the comparison groups (Table 1). Comparison groups were established reflecting *APOE3Ch* effect within the *PSEN1* E280A background for each individual patient, for both patients combined, and for *APOE3Ch* versus *APOE3* in a *PSEN1WT* background (Table 1). Statistical overrepresentation was performed using binomial test type and Bonferroni correction for four distinct differential gene expression groups. 75% of test groups came back positive for both Wnt signaling and Cadherin signaling with fold enrichment of 1.57 to 1.89 and 1.71 to 2.33 respectively. 25% of test groups came back positive for Heterotrimeric G protein signaling with fold enrichment of 1.76. (Table 1).

**Table 1:** PANTHER Pathways overrepresentation analysis. Differential expression gene lists were established using pseudo-bulk expression levels to the outlined comparison groups. PANTHER pathways was used to identify gross pathways influenced by *APOE3Ch* with and without the *PSEN1* background. Binomial test type and Bonferroni correction were used.

| Comparison group 1 | Comparison group 2 | PANTHER Pathways | Reference count | Input | Expected | Fold enrichment | Bonferroni P-value |
| --- | --- | --- | --- | --- | --- | --- | --- |
| ω- ApoE3Ch/PSEN1 E280A | ω- ApoE3/PSEN1 E280A | Blood coagulation (P00011) | 46 | 16 | 4.38 | 3.65 | 2.19E-03 |
|  |  | Cadherin signaling pathway (P00012) | 166 | 37 | 15.81 | 2.34 | 5.24E-04 |
|  |  | Wnt signaling pathway (P00057) | 314 | 51 | 29.91 | 1.71 | 3.79E-02 |
|  |  | Unclassified (UNCLASSIFIED) | 17971 | 1622 | 1711.65 | 0.95 | 0.00E+00 |
| α- ApoE3Ch/PSEN1 E280A | α- ApoE3/PSEN1 E280A | Cadherin signaling pathway (P00012) | 166 | 58 | 26.2 | 2.21 | 7.53E-06 |
|  |  | Alzheimer disease-presenilin pathway (P00004) | 131 | 39 | 20.67 | 1.89 | 3.02E-02 |
|  |  | Wnt signaling pathway (P00057) | 314 | 78 | 49.55 | 1.57 | 1.53E-02 |
|  |  | Unclassified (UNCLASSIFIED) | 17971 | 2734 | 2835.87 | 0.96 | 0.00E+00 |
| α- ApoE3Ch/PSEN1WT | ω- ApoE3/PSEN1WT | Cadherin signaling pathway (P00012) | 166 | 79 | 33.89 | 2.33 | 3.31E-09 |
|  |  | Wnt signaling pathway (P00057) | 314 | 119 | 64.1 | 1.86 | 6.09E-08 |
|  |  | Heterotrimeric G-protein signaling pathway-Gq alpha and Go alpha mediated pathway (P00027) | 125 | 45 | 25.52 | 1.76 | 4.48E-02 |
|  |  | Unclassified (UNCLASSIFIED) | 17971 | 3494 | 3668.57 | 0.95 | 0.00E+00 |
| α- ApoE3Ch/PSEN1 E280A | α- ApoE3/PSEN1 E280A | Unclassified (UNCLASSIFIED) | 17971 | 556 | 584.81 | 0.95 | 0.00E+00 |
| ω- ApoE3Ch/PSEN1 E280A | ω- ApoE3/PSEN1 E280A |  |  |  |  |  |  |

### β-catenin is elevated in APOE3 Christchurch cerebral organoids

We focused on characterization of rosettes, which are structures commonly observed in cerebral organoids resembling neural tubes including pseudostratified epithelium with apico-basal polarity (Di Lullo and Kriegstein 2017). Rosettes were smaller within *APOE3/PSEN1* E280A cell lines compared to their *PSEN1WT* or *APOE3Ch* counterparts, suggesting a differential maturation phenotype. Consistent with a more mature phenotype, *APOE3Ch* organoids stained prominently with *Reln*, a marker of more mature neurons (Figure 6 A-D) (Di Lullo and Kriegstein 2017; Lancaster et al. 2013). Canonical Wnt activation leads to accumulation of β-catenin and inhibition of GSK3β, a critical modulator of tau phosphorylation (De Ferrari et al. 2014; Jackson et al. 2002). Thus, we hypothesized Wnt/β-catenin/Cadherin pathway regulation could link *APOE* genotypes to tau phosphorylation via modulation of β-catenin. Distribution of β-catenin was prominent in apical regions close to the rosette’s lumen (ribbon-like) and more homogenously localized within the pseudostratified epithelium of organoids with *APOE3Ch* variant (Figure 6E). We segmented the body of the rosette and ribbon for quantification of β-catenin expression (Figure 6F-H). Rosette body and ribbon features show significant increase of β-catenin expression in *APOE3Ch* variant carrier organoids vs. control. This phenotype was not impacted by *PSEN1* genotypes suggesting a direct effect of the *APOE* genotype on tau phosphorylation phenotypes. We observed a marked decrease in rosette area within *APOE3/PSEN1* E280A organoids compared to *APOE3Ch* rosette size (Figure 6I). This observation did not carry into rosette aspect ratio (Figure 6J).

**Figure 6:**
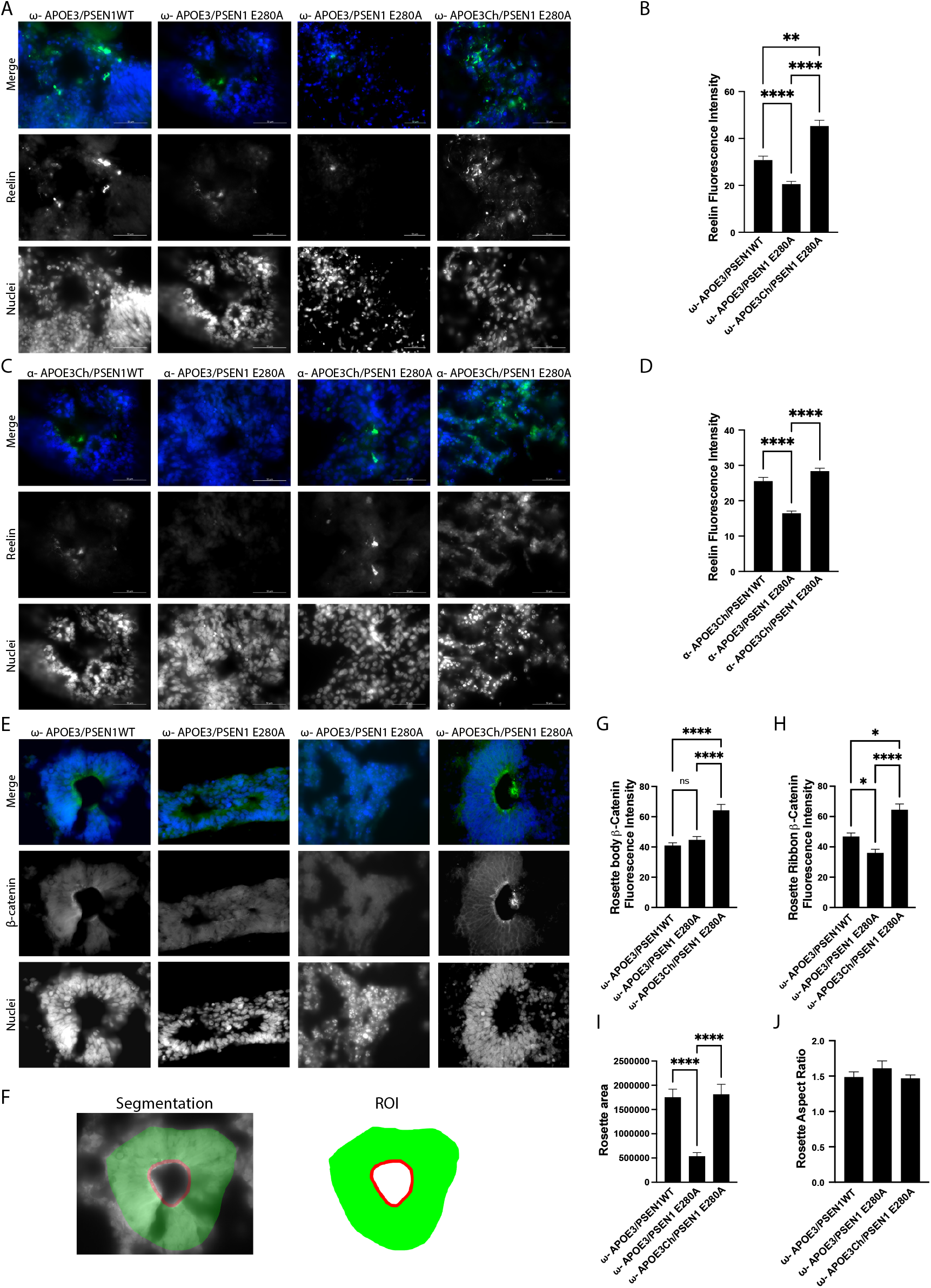
*APOE3Ch* influences Reelin, β-catenin, and neural rosette features. Immunofluorescence staining was performed on cerebral organoids to identify changes in protein of interest and physical data. Organoids were imaged at 63X and quantified. Representative images of Patient ω **(A)** and Patient α **(C)** stained for reelin and nuclei, scale bars 50 μm. Reelin channel intensity was averaged, with isogenic controls averaged together for Patient ω **(B)** and Patient α **(D)**. Cerebral organoids were stained for β-catenin, rosettes were identified in Patient ω and imaged at 63X **(E)**. Segmentation was performed and regions of interest defined **(F)**. β-catenin was quantified for both rosette body **(G)** and rosette ribbon **(H)**. Rosette physical features were then measured for area **(I)** and aspect ratio **(J).**

## Discussion

We have developed novel induced pluripotent stem cell lines derived from the ADAD Paisa kindred, used genetic engineering to remove the *PSEN1* E280A mutation as well as editing native *APOE3* to either add or remove the Christchurch mutation, formed cerebral organoids, identified pathways through RNA-seq, and supported these findings through immunostaining. While it is well documented that iPS cell systems can model aspects of Alzheimer’s disease pathology (Penney, Ralvenius, and Tsai 2020), we have demonstrated that the iPS cell system is capable of modeling disease resistance through specific patterns of tau posttranslational phosphorylation: protective T205 site (Ittner et al. 2016) and detrimental S202/T205 and S396 sites (Mondragón-Rodríguez et al. 2014). We have found that the *APOE3Ch* variant alters the translational landscape to promote changes in Cadherin and Wnt signaling, which affects β-catenin, irrespective of the *PSEN1* background. Taken together, our data suggests that targeting processes affected outside of *PSEN1*/Amyloidβ/Tau pathways may be therapeutically viable in the treatment of Alzheimer’s disease, and potentially other tauopathies including fronto-temporal dementia.

*PSEN1* mutations may impact neuronal differentiation in iPS cells derived systems, such as cerebral organoids. This effect appears to be mediated by dysfunctional gamma secretase activity leading to reduced Notch signaling in stem cells and premature aging phenotypes (Arber et al. 2021). We noticed some developmental differences in the organoids carrying the *PSEN1* E280A genotype. However, our cultures were relatively young, preventing the development of a fully senescent neuronal phenotype. On the other hand, it is remarkable that even at a relatively young developmental stage, organoids with the *PSEN1* E280A genotype can show already abnormal tau phosphorylation as identified by S202/T205 and S396 phosphorylations. This shows that young cerebral organoids are already valid as pathological models for AD and for possible mechanisms of protection.

The effects of *APOE3Ch* on Wnt and Cadherin signaling uncovered by scRNA-sequencing of cerebral organoids were unexpected and may operate via multiple mechanisms ultimately resulting in β-catenin upregulation. Cadherins are a family of calcium dependent transmembrane adhesion proteins that link β and α catenin to the actin cytoskeletal network (Punovuori, Malaguti, and Lowell 2021) and also regulate cellular homeostasis through signaling mediating development, proliferation, apoptosis, and disease pathology (Yulis, Kusters, and Nusrat 2018). Cadherins regulate calcium dependent cell-cell adherent junctions, where the chelation of calcium abolishes adhesive activity and allows proteolytic degradation of cadherins (Nagar and Overduint 1996; Kim et al. 2011). Thus, proper calcium levels play a vital role in cellcell dynamics as well as maintaining a pool of cadherin.

Wnt signaling can be assigned into two pathways, canonical Wnt signaling or the non-canonical planar cell polarity (PCP) and Wnt/Calcium pathway subdivisions. Canonical Wnt signaling requires extracellular Wnt binding to LRP5/6 and Frizzled for signal transduction across the cell membrane to Disheveled. Once internalized, the signal is passed to the β-catenin destruction complex, a proteinaceous structure composed of GSK3β and other proteins, resulting in the release of β-catenin by inhibition of GSK3β. Free β-catenin is then able to translocate to the nucleus and activate TCF/LEF transcription. In the PCP pathway, Wnt directly binds to Frizzled and transduces the signal to Disheveled, which in turn activates RhoA and Rac1 and eventual JNK pathways. In the non-canonical calcium dependent subpathway, Wnt binds directly to Frizzled, transduces the signal to Disheveled and interacts with trimeric G proteins and phospholipase C, increasing intracellular calcium concentration inducing CamKII and calcineurin activation (Inestrosa and Varela-Nallar 2014). CaMKII is vital in controlling NMDA receptor activity (Incontro et al. 2018), which also acts as a calcium channel (Lau et al. 2009).

GSK3β is a protein kinase that phosphorylates and primes tau for inclusion in paired helical filaments and fibrils (Hooper, Killick, and Lovestone 2008). Indeed, GSK3β is known to phosphorylate tau at the early pathology sites S396 and at S202 (Li and Paudel 2006). GSK3β is also a vital component of the β-catenin destruction complex. It is responsible for phosphorylating β-catenin for ubiquitination and proteosomal degradation (Inestrosa and Varela-Nallar 2014). This persistent degradation maintains low levels of free cytoplasmic β-catenin and inhibits gene transcription. Cerebral GSK3β stimulation by phosphorylation at Y216 is mediated by intracellular calcium levels and by calcium-dependent PYK2 (Hartigan and Johnson 1999; Hartigan, Xiong, and Johnson 2001; Sayas et al. 2006).

*APOE* is the most significant, known, risk factor for sporadic Alzheimer’s disease, ApoE4 exhibits the strongest receptor binding and is considered a high risk allele, while ApoE2 exhibits the weakest receptor binding and is considered protective (Yamazaki et al. 2019). The *APOE3Ch* variant was found in a protected ADAD subject and was shown to have weaker binding than its ApoE3 counterpart to heparin sulphate proteoglycans (Arboleda-Velasquez et al. 2019), however the pathway of protection was unknown. Imputation of genetic causality was also not feasible because of the rarity of the Christchurch variant. Thus, the need for genetic analyses *ex vivo* as conducted here.

In sum, our data suggests that iPS-derived cerebral organoids can be informative of protective mechanisms, and they show experimentally for the first time that the Christchurch variant could be mechanistically linked to a protective pattern of tau phosphorylation. Importantly our data showed a prominent role for Wnt and Cadherin signaling in the presence of the *APOECh* variant. β-catenin is differentially regulated in *APOE3Ch* cerebral organoids, which affects Wnt/Cadherin signaling and GSK3β activity. This finding may open the door for novel therapeutics focused on *APOE3Ch* and the regulation of Wnt, Cadherin, and β-catenin.

## Acknowledgements

The authors thank the Colombian families with ADAD for their contributions and extraordinary commitment to research. We would like to thank L. Daheron and the Harvard Stem Cell Institute for their effort and support. We also thank A. Koutoulas for technical support with genome sequencing, K. Hartmann and C. Dwumfour for technical support with immunohistochemical staining, and S. Levine and S. Mildrum of the BMC core at the Massachusetts Institute of Technology.

## Data Availability

Anonymized clinical and imaging data are available upon request, subject to an internal review by F.L., and Y.T.Q. to ensure that the participants’ confidentiality, completion of a data sharing agreement, and in accordance with University of Antioquia’s and Massachusetts General Hospital’s IRB and institutional guidelines. Experimental data is available upon request, subject to Massachusetts General Hospital and Schepens Eye Research Institute of Mass Eye and Ear institutional guidelines. iPS cells requests will be considered based on a proposal review, completion of a material transfer agreement and/or a data use agreement, and in accordance with the Neuroscience Group of Antioquia institutional guidelines upon peer reviewed publication. Please submit requests for participant-related clinical and imaging data and samples to Y.T.Q.; requests for experimental data and single-cell RNA sequencing data to J.F.A.-V.; and, requests for iPS cells to F.L.

## Methods

### Patient selection and sample collection

We have selected two patients for this study, they will be known in this study as Patient α and Patient ω to ensure patient privacy. Patient α was previously described as being a protected patient from familial Alzheimer’s disease (Arboleda-Velasquez et al. 2019). Patient α was part of the Paisa *PSEN1* E280A kindred in her seventies at time of mild cognitive impairment. She was found to have the *APOE3* R136S Christchurch variant which provided her resistance to Alzheimer’s development. Patient ω was also selected as a Paisa kindred female with development of ADAD at expected age of onset and which PET imaging data was available with expected brain pathology.

Blood samples from each individual were obtained by venipuncture. Peripheral blood mononuclear cells were separated by Ficoll-Hypaque 1077 and submitted for reprogramming and genetic editing.

### *In vivo* neuroimaging

Structural magnetic resonance imaging (MRI), Pittsburgh compound B (PiB) and Flortaucipir (FTP) positron emission tomography (PET) were performed at Massachusetts General Hospital as described elsewhere (Quiroz et al. 2018). Briefly, MRI images were processed with FreeSurfer (FS, v 6.0) to identify surface boundaries and standard regions of interest (Desikan et al. 2009). PET data were acquired and processed according to previously published protocols (Johnson et al. 2016), whereby PiB data were expressed as distribution volume ratios (DVR, Logan, 0-60 min.) and FTP as standardized uptake value ratios (SUVR, 80-100 min.), both using cerebellar gray matter as the reference region. PET images were affine co-registered to each subject’s T1 images and visualized using FS surface projections (sampled at gray matter midpoint, surface-smoothed 8mm). No partial volume correction was applied to PET images for the purposes of this study.

### Reprogramming and genetic editing

Cell services were performed using the Harvard Stem Cell Core: whole blood was reprograming though Cytotune 2.0 (ThermoFisher), colonies were allowed to grow and were then assessed for the iPS markers SSEA, Oct4, Tra-1-60, and Nanog using immunocytochemistry as well as qPCR for trilineage. Successful screened colonies were then processed further for genetic editing. Guide RNA (gRNA) and single stranded oligodeoxynucleotide (ssODN) sequences were determined (Supplementary Table 1) and colonies were karyotyped.

### Cell culture

Reprogrammed and edited cells were grown on hESC-qualified (human Embryonic Stem Cell) Matrigel (Corning #354277) using mTeSR Plus (StemCell Technologies #100-0276) supplemented with the antimicrobial Normocin (Invivogen #ant-nr-1) and clump passaged weekly using ReLeSR (StemCell Technologies #05872) according to manufacturers recommended protocols. Cell clumps were sparsely plated to allow for easy physical removal of spontaneously differentiated cells. Cells were grown at 37°C and 5% CO_2_.

### Differentiation

Cells were grown as described above. Regions of spontaneously differentiated cells were identified and physically removed. Organoids were made using StemDiff Cerebral Organoid Kit (StemCell Technologies #08570) and Normocin using manufacturers recommended protocol, however iPS spheroids were made using EB formation media (Stem Cell Technologies #05893) with Normocin. Briefly, cells were detached to single cell using Accutase (StemCell Technologies #07920) and counted (Countess II) using Trypan Blue. 9,000 cells/well were plated into low retention 96well U-bottom plates (S-Bio #MS-9096UZ) in EB formation media with γ-27632 (ATCC #ACS-3030) with regular fresh media additions. After 5 days, spheroids were transferred to a 24well flat bottom low retention plate (Corning #3473) to induce differentiation. Organoids were then embedded into individual hESC-qualified Matrigel droplets and plated into 6-well flat bottom low retention plates (Corning #3471) for expansion. Organoids were then placed on to a shaker plate inside the incubator with regular media exchanges (2-3 days) until downstream analysis.

### Immunostaining

Organoids were fixed in 4% PFA for 2 hours then washed in 1xdPBS thrice. Organoids for IHC analyses were embedded in paraffin, and sectioned at 2 μm. After dewaxing and inactivation of endogenous peroxidases (PBS/3% hydrogen peroxide), antibody specific antigen retrieval was performed using the Ventana Benchmark XT machine (Ventana, Tuscon, Arizona, USA). Sections were blocked and afterwards incubated with the following primary antibodies: anti-GFAP (M761, DAKO GmbH), IBA1 (NC9288364, Wako Chemicals), PAX6 (AB_528427, DSHB, we thank the DSHB for the usage of this PAX6 antibody), OLIG2 (AB9610, Millipore), APOE (Mab947, Millipore), MAP2 (13-1500, ThermoFisher), TAU (AHB0042, ThermoFisher), pTAU S202/T205 (AT8, MN1020, ThermoFisher), pTAU T205 (AB254410, Abcam), and KI67 (275R-15, Cell Marque). For detection of specific binding, the Ultra View Universal 3,3’-Diaminobenzidine (DAB) Detection Kit (760-500, Ventana, Roche) was used, which contains secondary antibodies, DAB stain and counter staining reagent for detection of nuclei. For detection of APOE antibody binding, the anti-goat Histofine Simlpe Stain MAX PO immune-enzyme polymer (medac GmbH, 414161F) was used. Staining were evaluated by an experience pathologist and representative images were taken with a Leica DMD108 digital microscope.

Organoids for immunofluorescence analyses were infiltrated with 30% sucrose until they dropped and frozen in OCT and sectioned at 15 μm. Sections were washed with 1xdPBS and blocked for 1 hour (PBS, 3% BSA, 0.2% Triton X-100 and 0.02% azide). Then, they were incubated in primary antibody diluted in blocking buffer using the following primary antibodies: anti-Phospho-Tau (Ser396) (1:500, 44-752G, Invitrogen), anti β-catenin (E-5) (1:50, sc-7963, Santa Cruz) and anti-Reelin (CR-50) (D223-3, MBL Life Science) overnight at 4°C. After washing the sections with 1xdPBS, they were incubated in the following secondary antibodies: Donkey anti-rabbit IgG Alexa fluor 647 (1:500, A-31573, ThermoFisher) and Donkey anti mouse IgG Alexa fluor 488 (1:500, A-21202, ThermoFisher). DAPI solution was used for nuclei staining. Images were taken at 10X and 63X magnification using a ZEISS Axioscope digital microscope.

### scRNA-seq

Six organoids from each cell line were collected and processed using MACS papain neural tissue dissociation (Miltenyi Biotec #130-092-628) according to manufacturer’s protocol and resuspended in 0.22μm filter sterilized dPBS 0.04%BSA solution. Cells were serially filtered through a 70μm (Miltenyi Biotec #130-110-916) then 40μm (Bel-Art #H136800040) strainer to remove clumped cells. Samples were kept on wet ice and assessed at the BioMicroCenter core facility (Massachusetts Institute of Technology) for viability, cell density, and quality. Samples were processed on 10X Genomics Chromium Controller at the BMC Core facility. Sequences were then processed through the 10X Genomics Cell Ranger Suite.

### Transcriptomic analysis

The h5 files were read with the Seurat Read10X_h5() function (Seurat (Hao et al. 2021), an R package for scRNA-seq clustering and integration). The DoubletFinder (McGinnis, Murrow, and Gartner 2019) doubletFinder_v3() function was employed to identify and remove likely multiplets, the predicted multiplet rate was 0.8% per “Targeted Cell Recovery” of 1000 cells (Chromium). Sample subsets of 6500 cells were processed via DietSeurat() (default values) and saved as RDS files.

Sample integration was accomplished by merging Seurat objects with merge(), v.1 SCTransform() (Hafemeister and Satija 2019), and Harmony integration [RunHarmony()] (Korsunsky et al. 2019). Seurat objects were saved as RDS files.

Aggregate gene expression [AggregateExpression()] in clusters was employed to compare gene expression between samples. Aggregate expression values were used to generate tables with aggregate sums and log2 adjusted sample ratios.

Comparison groups were established (Table 1) reflecting log2(fc >1 and <-1) and pathway analysis run using PANTHER Pathways Overrepresentation Test version 17.0 released 2022-02-22 with *Homo sapiens* (all genes in database) reference genome using a Binomial test type and Bonferroni correction.

Gene Set Enrichment Analysis (GSEA) (Subramanian et al. 2005) was performed using fgsea to facilitate cell type identification. The MSigDB cell type signature (C8) gene set database (https://www.gsea-msigdb.org/gsea/msigdb/index.jsp, v7.4) was queried (the C8 gene set was filtered to retain only brain-relevant entries)(Cao et al. 2020; Zhong et al. 2018; La Manno et al. 2016; Fan et al. 2018). NES (enrichment score normalized to mean enrichment of random samples of the same size) scores >= 7.5 were considered significant and informed cell type assignment. Clusters with NES scores < 7.5 were designated “unidentified” (see Supplemental Files). Clusters were further classified by general cell type into five “superclusters”. These are “neuronal”, “glial”, “progenitor”, “pericyte”, and “undetermined” (Figure 5C).

### Immunofluorescence image quantification

Rosette β-catenin quantification was implemented in MATLAB 2021a with use of the Image Processing Toolbox. We quantified the average channel intensity in two types of rosette ROIs (regions of interest): the Body and the Ribbon. The masks (or binary image representations) of the Body and Ribbon were produced via a combination of hand-drawing (to get the outer boundary and inner luminal boundary lines) and Otsu thresholding to remove background pixels from the hand-drawn regions. The hand-drawn mask for the rosette Body was defined as the region between the inner and outer boundaries eroded by 1.8 μm (to ensure that the ribbon and outer edges were not included in the Body ROI). The rosette Ribbon ROI was defined as the inner luminal boundary line dilated by 1.8 μm to approximately capture the full ribbon thickness.

Rosette area was measured by summing the number of pixels within the rosette body and ribbon ROIs. The rosette Aspect Ratio was defined as the major axis length divided by the minor axis length. We used MATLAB function *regionprops* to measure the major and minor axis lengths.

In both pTau S396 and Reelin fluorescence imaging, background intensity was subtracted via Top-hat filter to remove non-uniform background illumination. Top-hat filtering was accomplished with a large structuring element (54 μm radius disk) so as not to remove signal from large features of interest. After background removal we used Otsu’s threshold method to find the foreground and computed the mean intensity over the foreground.

---

**Supplemental table 1:**
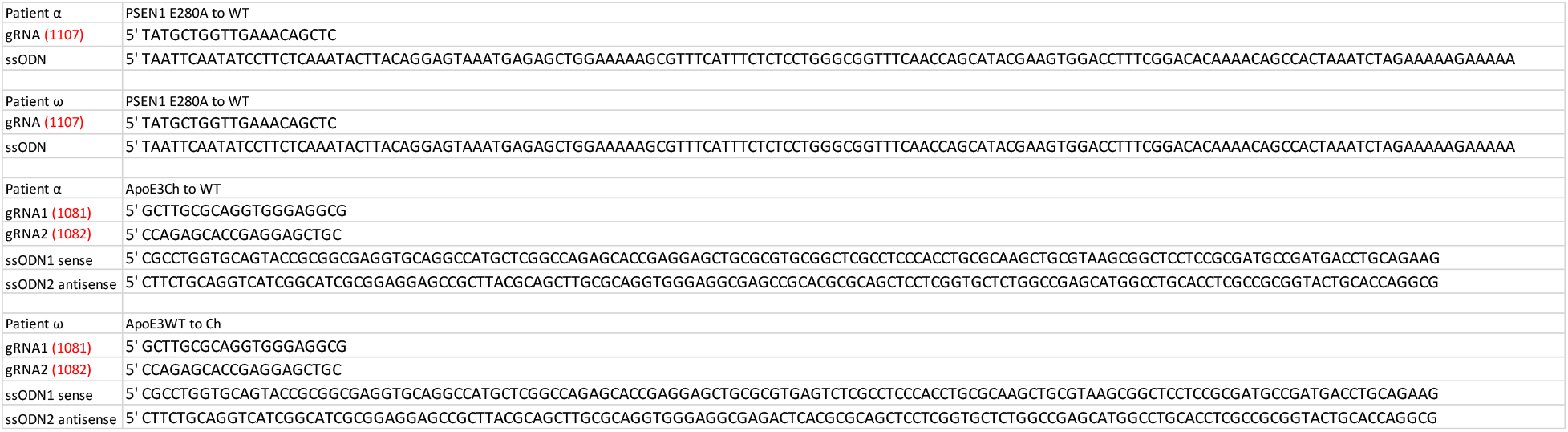
Primer sequences used for genetic editing of patient cell lines.

